# Superfast periodicities in distress vocalizations emitted by bats

**DOI:** 10.1101/734640

**Authors:** Julio C. Hechavarría, M. Jerome Beetz, Francisco Garcia-Rosales, Manfred Kössl

## Abstract

Communication sounds are ubiquitous in the animal kingdom, where they play a role in advertising physiological states and/or socio-contextual scenarios. Distress sounds, for example, are typically uttered in distressful scenarios such as agonistic interactions. Here, we report on the occurrence of superfast temporal periodicities in distress calls emitted by bats (species *Carollia perspicillata*). Distress vocalizations uttered by this bat species are temporally modulated at frequencies close to 1.7 kHz, that is, ∼17 times faster than modulation rates observed in human screams. Fast temporal periodicities are represented in the bats’ brain by means of frequency following responses, and temporally periodic sounds are more effective in boosting the heart rate of awake bats than their demodulated versions. Altogether, our data suggest that bats, an animal group classically regarded as ultrasonic, can exploit the low frequency portion of the soundscape during distress calling to create spectro-temporally complex, arousing sounds.

## Introduction

Communication calls are ubiquitous in the animal kingdom, where they play a role in advertising physiological states and socio-contextual scenarios ^1–4^. Several types of communication calls have been described according to the context in which they are produced. Fearful screams, for example, are used in distressful scenarios such as agonistic interactions ^1,5^. On the other hand, positive affect vocalizations are broadcasted in pleasuring situations, such as tickling ^6,7^, while mating and isolation vocalizations are produced in sexual contexts ^8,9^ and in the context of mother-infant interactions ^10–12^, respectively.

Bioacoustics studies have shown that different animal species rely on different spectro-temporal repertoires for advertising socio-contextual cues. Yet, despite the large vocalization variability found throughout the animal kingdom, rules have been described that can explain the physical attributes of sounds broadcasted by different species in different contexts. For example, “motivational-structural rules” proposed by Morton^13^ predict large differences in spectro- temporal designs between vocalizations emitted in appeasing and agonistic contexts. While agonistic vocalizations tend to be broadband and harsh in their spectrum, appeasing vocalizations tend to be pure-tone like and have low harmonic-to-noise ratios. Such differences between agonistic and non-agonistic utterances have been well documented in a wealth of studies in vertebrates ^1,14–21^.

Studies on the human soundscape have reported that differences between agonistic and non-agonistic sounds also exist at the level of temporal modulations. For example, in humans, recognizing sounds as screams is linked to the sounds’ temporal periodicities that appear in the form of “roughness”, e.g. amplitude modulations (AMs) at frequencies close to 100 Hz ^1^. In humans, in addition to naturalistic screaming, rough sounds also are used to express pain, suffering and aggressiveness in musical compositions such as the opera and hard rock ^22,23^, as well as for communicating urgency in artificial alarm systems ^1^. Besides humans, rough/periodic sounds have also been reported in vocalizations of other animal groups, including primates, otters and birds ^24–29^, although in many of these species, roughness (and fast periodicities in general) have not been studied quantitatively using methods similar to those employed for characterizing the human soundscape.

The main aim of this study is to test whether fast periodicities are a distinctive feature of distress calls emitted by bats, a highly vocal animal group. Bats use acoustic signals for a dual purpose, namely, echolocation and communication. The former is important for orientation in the dark while the latter is important for coordinating complex social behaviors ^4,30,31^. A type of communication vocalizations uttered by bats are the so-called distress signals, which are produced when the animals are tangled in catch nets or caught by a predator or a person ^32–35^. Bat distress calls are typically noisy and broadband, in accordance with Morton motivational structural rules ^32,35^, and they trigger exploratory and mobbing behaviors in individuals from the same and other species ^33,34,36,37^. Bat distress calls are also known to trigger neural responses in the amygdala ^38^, to entrain field potentials and spiking in the auditory cortex ^39–41^, and to boost activity in the hypothalamic-pituitary axes ^42,43^.

In this article, we studied the amplitude modulation pattern of bat distress vocalizations and searched for fast periodicities. Our hypothesis was that bat distress utterances should carry fast amplitude fluctuations. This hypothesis was based on the idea that fast amplitude fluctuations (occurring in the form of roughness in humans) could be a generalized trait for signaling distress in the vertebrate clade. The data corroborated this hypothesis. In bats, fast periodicities occur linked to a low fundamental frequency (a putative acoustic correlate of pitch) that lies outside the range of audible frequencies for the bat species studied. This result is important and novel as it highlights that bats, an animal group classically regarded as ultrasonic, can exploit the low frequency soundscape to create rhythmic amplitude fluctuations in the uttered sounds. In addition, we show that fast periodicities are represented in the bats’ brain by means of frequency-following responses, and that listening to vocalizations containing fast periodicities boosts the heart rate (HR) of conspecific bats. These results suggest that fast-periodic sounds might indeed play a role in signaling distressful contexts in bats.

## Results

### Bat distress syllables carry superfast periodicities

We recorded distress calls from 13 adult bats (6 females and 7 males) of the species *Carollia perspicillata*. This species emits sequences of distress calls composed of basic vocalization units defined as “syllables” ^35^. In bats, the production of distress calls can be triggered by holding the animals in the hands while carefully caressing the neck-skin ^33,34^. All procedures used in this article to obtain distress calls in bats comply with current laws for animal experimentation in Germany (Regierungspräsidium Darmstadt, permit # F104/57) and were optimized for recording bat vocalizations for time periods no longer than 3 minutes.

We studied a total of 114 distress “sequences”. Each of those sequences was composed of sound units defined as “syllables” (see also ref. ^35^ for the definition of distress sequences and syllables). An example distress sequence is shown in figure 1A. This particular sequence contained 71 syllables arranged over a time period of 2.38 s. As shown in figure 1A and in a previous article^35^, within a distress sequence, syllables are temporally arranged in groups defined as “multi-syllabic bouts”. A zoom-in into the multi-syllabic bout containing syllables 55-59 is shown at the bottom of figure 1A to illustrate the temporal separation between syllables.

**Figure 1.**
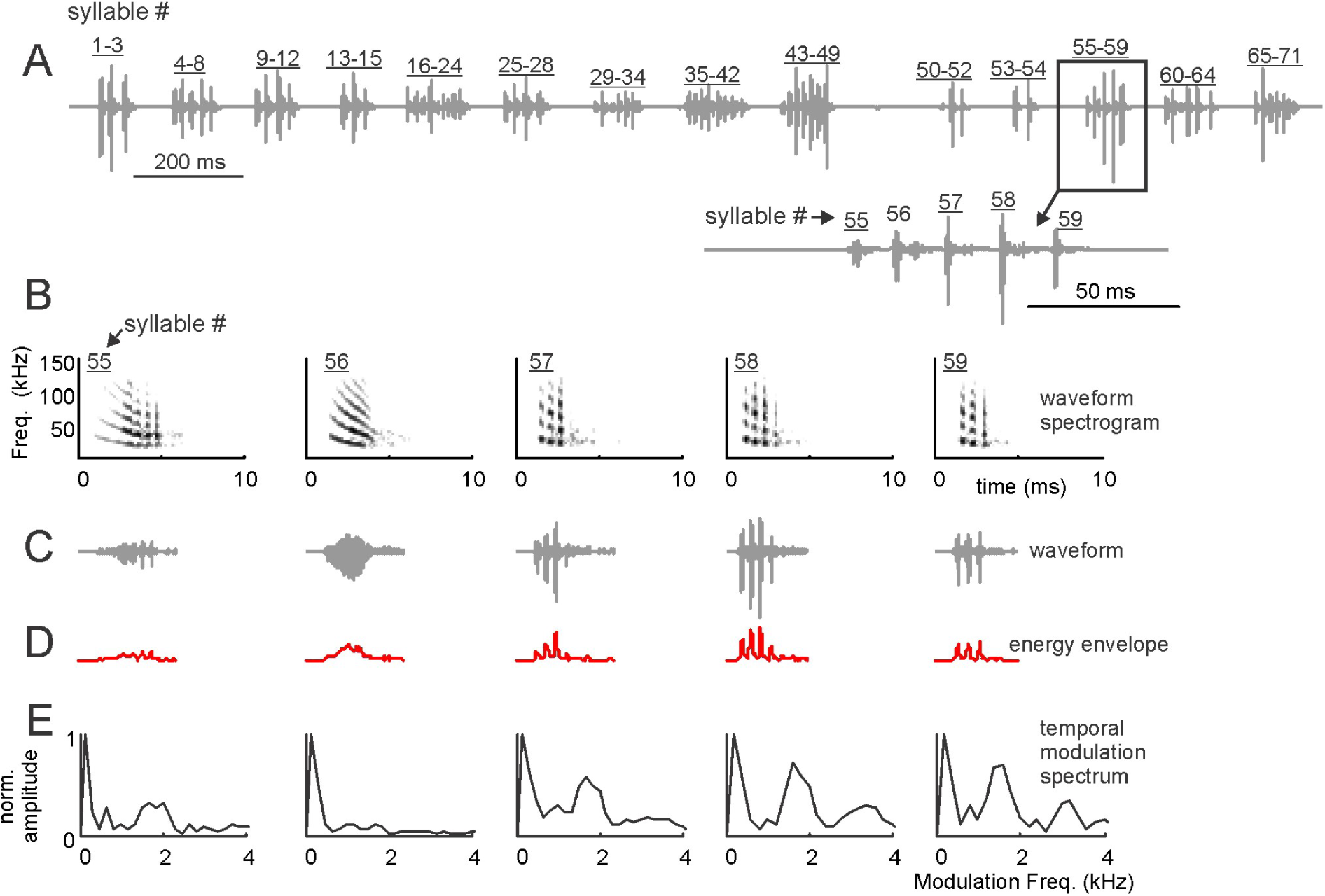
Bat distress vocalizations carry superfast periodicities. (A) Example distress sequence containing 14 syllable groups (bouts) and 71 syllables. A zoom-in into the bout composed of syllables 55-59 is provided. (B) spectrograms, (C) time waveforms, (D) envelope and (E) temporal modulation spectra (TMS) of syllables 55-59. Note that fast periodicities at ∼ 1.7 kHz occur in syllables 55, 57-59, but are less clear in syllable 56.

We searched for fast periodic amplitude fluctuations in individual distress syllables. The procedure used for analyzing single syllables is illustrated in figure 1B-E for the syllables that formed the multi-syllabic bout represented in the bottom panel of figure 1A. The energy envelope of each syllable was calculated (Fig.1D) and the spectrum of the envelope (defined as the temporal modulation spectrum, TMS, Fig. 1E) was obtained and analyzed in the range between 0-4 kHz. Note that this range contains frequencies that are well above temporal modulation frequencies found in rough vocalizations of humans (i.e. ∼ 100 Hz ^1^). Analyzing such broad range of temporal modulation frequencies was important, since we did not know the frequency range at which fast periodicities could appear in bat vocalizations.

As it can be noted in the example distress syllables represented in figure 1B-E, a single distress sequence could contain syllables with different types of TMS. For example, the TMS of syllables 55, 57, 58 and 59 (Fig. 1B-E) had a pronounced peak at ∼ 1.7 kHz. In syllable 56, the peak at ∼1.7 kHz was less evident. We reasoned that syllables modulated at rates ∼ 1.7 kHz could be classified as fast periodic vocalizations (FPVs) since they contained a pronounced temporal modulation pattern. Note that such pronounced signal periodicity can indicate the fundamental frequency of the syllables and can thus be interpreted as an acoustic correlate of signal “pitch” and/or as acoustic “non-linearities” (see below and discussion). Also note that 1.7 kHz is a rather low frequency for *C. perspicillata*, a bat species that can reach frequencies above 100 kHz both while echolocating and while producing communication calls ^35,44^. In fact, the cochlear frequency response curves of *C. perspicillata*, calculated using distortion product otoacoustic emissions, suggest that hearing in this animal species deteriorates at frequencies below 5 kHz (see upcoming text in the results section).

We classified the distress syllables recorded into FPVs or non-FPVs based on their TMS. For that purpose, we relied on a binary support vector machine (SVM) classification algorithm that was fed with the TMS of all 7025 syllables recorded. The SVM classifier was trained with two sets of TMS comprising the TMS of 50 FPVs and 50 non-FPVs. The training TMS sets are shown in supplementary Figure S1 and they were chosen based on visual inspection of the TMS of randomly chosen syllables. Only syllables containing pronounced peaks in the 1.15-2.45 kHz range were selected for the FPV training set, and only syllables that contained no clear energy peaks in that range were selected for the non-FPV set.

The results obtained with the SVM classifier are depicted in figure 2. Altogether, 3349 out of 7025 syllables studied (47.7%) were classified as FPVs. The TMS of all FPVs and non-FPVs is shown as a colormap in Fig. 2A and B, respectively. Note that in the range from 1.15 to 2.45 kHz, brighter colors are present in the population of FPVs when compared to non-FPVs. This range is marked by a rectangle in figure 2A-B and it was defined as the “Frequencies of Interest” (FOIs) for further analysis. The presence of high energy at the FOIs was also visible in median curves for the populations of FPVs and non-FPVs (Fig. 2C and D) identified by the SVM classifier. To validate periodicity differences at the population level, we calculated the area under the curve at the FOIs in the two syllable groups (Fig. 2E). As expected, the power at the FOIs was significantly higher in FPVs than in non-FPVs (Wilcoxon ranksum test, p< 10^-200^). We also calculated the Cliff’s delta (*d*) to assess the size effect of FOI power comparisons. Briefly, *d* can take values between −1 and 1 and it measures how often values in one distribution are larger than values in a second distribution. Note that size effect calculations complement null-hypothesis testing and they are useful for ascertaining the meaningfulness of statistical significance obtained specially in the presence of large sample sizes (in our case n= 3349 and n= 3676 for FPVs and non-FPVs, respectively). The *d* value obtained from the comparison of FOI power in FPVs and non-FPVs was 0.94, indicating a large size effect (negligible effect: absolute *d* value (abs (*d*)) < 0.147, small: 0.147 <abs(*d*)<0.33, medium: 0.33<abs(*d*)<0.474, large: abs(*d*)>0.474, values after^45^).

**Figure 2.**
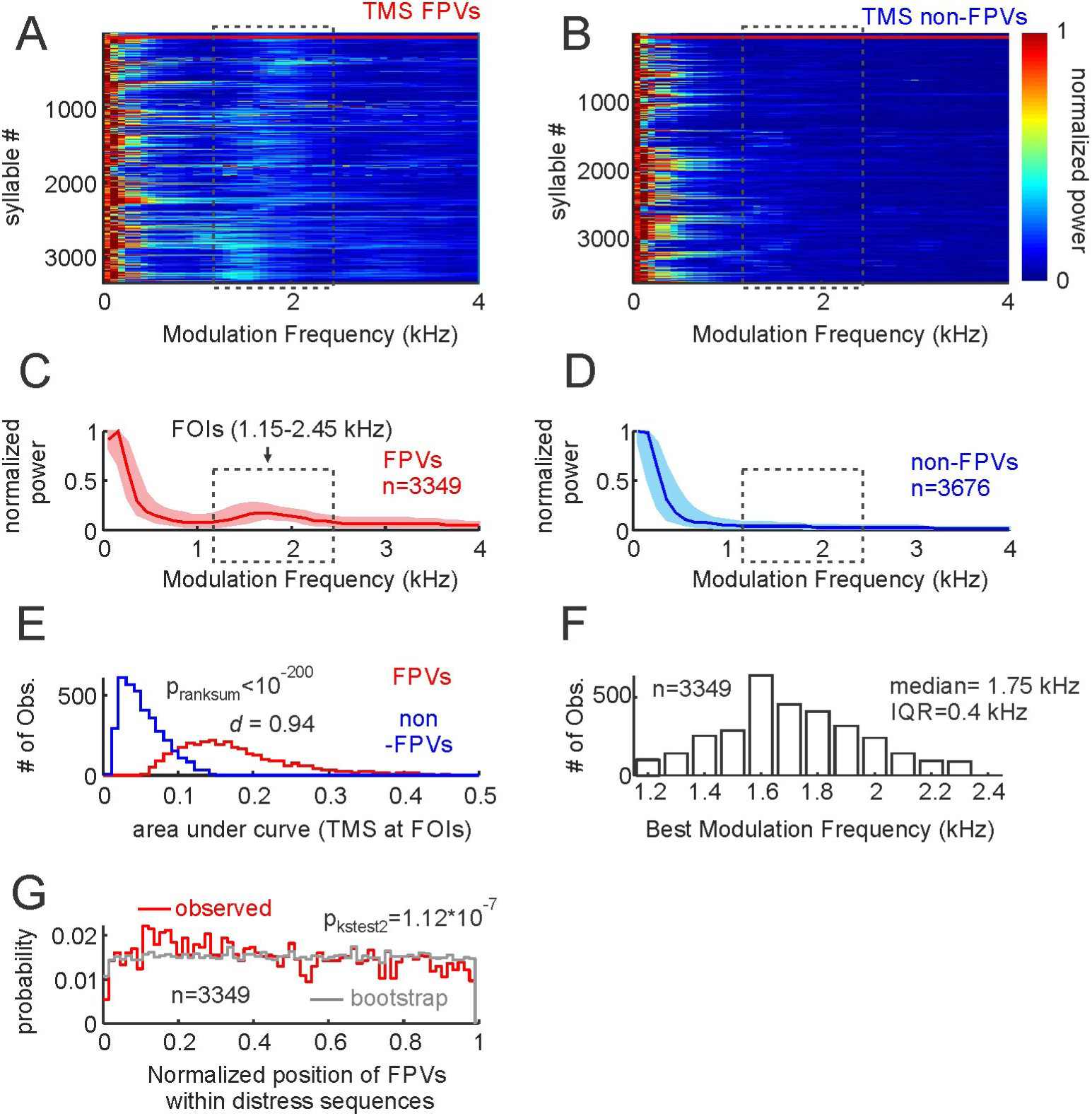
Temporal properties of fast periodic vocalizations (FPVs) and non-FPVs. A and B show the TMS of the two syllable groups, represented as colormaps. The first 50 FPVs and non- FPVs in the colormaps (border marked by horizonal red lines) were used to train the support vector machine classifier (see also supplementary figure S1). C and D are median TMS of all FPVs and non-FPVs studied (25th and 75th percentiles shown as shaded areas). Note that a peak occurs at the frequencies of interest (FOIs) in FPVs but not in non-FPVs. (E) Histogram of the area under the TMS curve at the FOIs in FPVs and non-FPVs. The p-value of a Wilcoxon ranksum test and the d size-effect metric are provided. (F) histogram of best modulation frequencies found in the population of FPVs. Median and inter-quartile range are provided. (G) Probability of finding FPVs in certain positions along the distress sequences. To calculate probability values, the relative position of each FPV was obtained taking into account the length of the sequence in which it occurred. This “observed” probability distribution was compared (Kolmogorov-Smirnov two-sample test) with an “expected” distribution obtained by randomly swapping the position of syllables within each sequence 100 times.

The best modulation frequency (BMF) of each FPV was calculated by searching for the frequency that contained the highest energy in the FOI range (that is, between 1.15 and 2.45 kHz). The BMF distribution had a median of 1.75 kHz with an interquartile range (IQR) of 0.4 kHz (Fig. 2F). We determined if FPVs occurred at a preferred position within distress sequences. To that end, we calculated the normalized position of each FPV relative to the length of the sequence in which it occurred. Though FPVs occurred throughout the sequences, the distribution of preferred positions was slightly skewed to the left (Fig. 2G, median =0.45, IQR= 0.49). The latter points towards a higher probability of finding fast periodic syllables in the first half of the sequences, independently of sequence length. This trend was validated statistically by comparing the temporal syllable distribution observed to a bootstrap distribution created by randomizing the positions of FPVs and non-FPVs in each sequence (100 randomizations for each sequence, two-sample Kolmogorov-Smirnov test: p = 1.12*10^-7^, Fig. 2G). Comparing the probability of finding FPVs in the first and second sequence halves also indicated statistical significance (Signtest, p=0.03).

### FPVs and non-FPVs differ in their spectral properties

Differences between FPVs and non-FPVs regarding their spectral properties were also tested. At the population level, there was a tendency for FPVs to have a narrower spectrum than non-FPVs, with non-FPVs tending to have higher power in the range from 40-80 kHz. The latter is visible in both the normalized spectra of all FPVs and non- FPVs (colormaps in Fig. 3A, B) and in the median spectra of the two syllable groups (Fig. 3C, D). Differences in spectral bandwidth between the two syllable groups were statistically significant, as validated by a ranksum test that compared the area under the normalized spectra in FPVs and non-FPVs (Fig. 3E, p_ranksum_<10^-121^). Note that the *d*-metric obtained for this comparison indicated a medium/small size effect (*d*=0.33). Besides these small differences in spectral bandwidth, FPVs and non- FPVs also differed in their peak frequencies (Fig. 3F, median_FPVs_=22 kHz, median_non___FPVs_=27 kHz, p_ranksum_=10^-121^) with FPVs tending to have lower peak frequency values, although the size effect in this case was also small (*d*=0.32).

**Figure 3.**
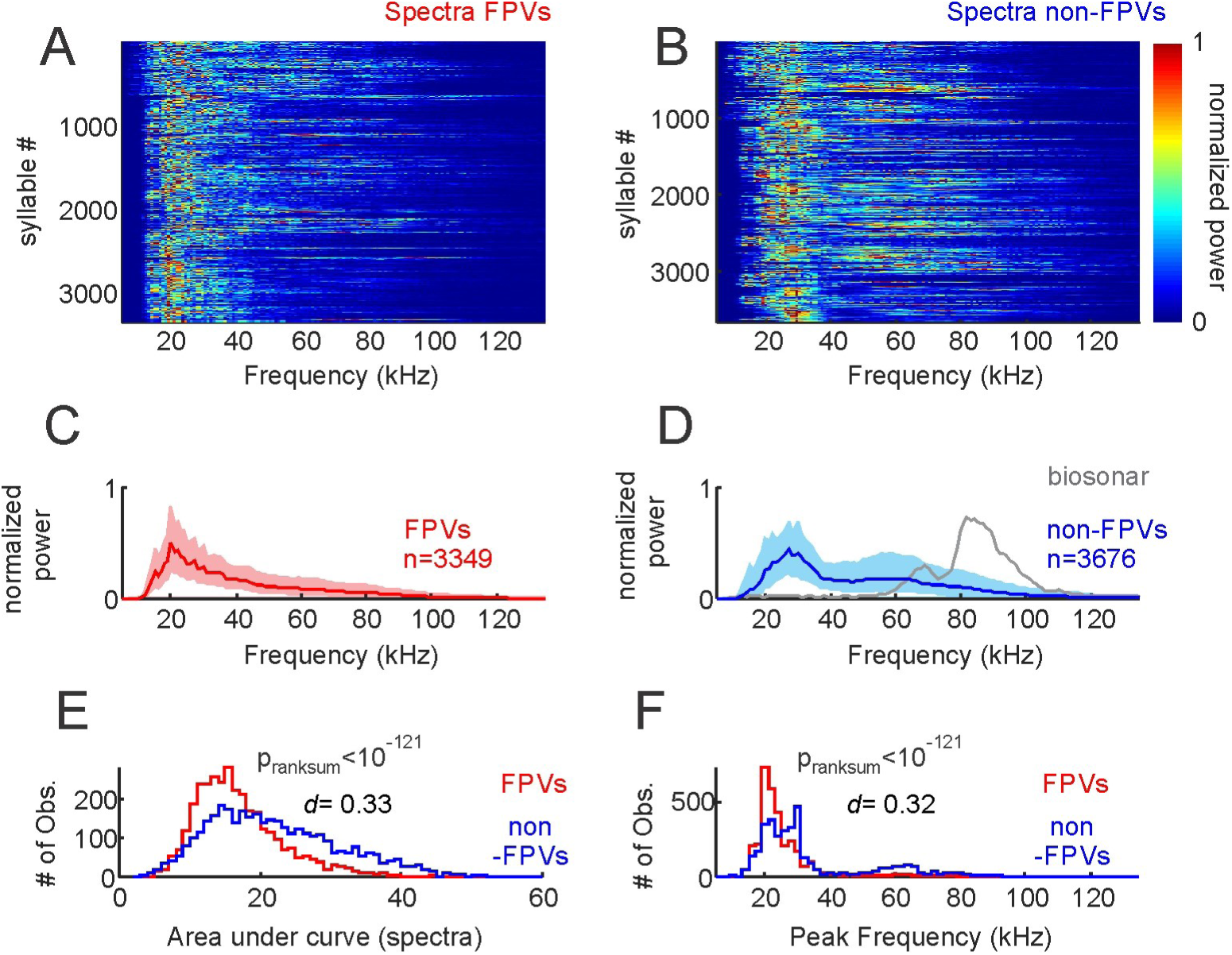
Spectral properties of fast periodic vocalizations (FPVs) and non-FPVs. A and B show the spectra of the two syllable groups, represented as colormaps. C and D are median spectra of all FPVs and non-FPVs studied (25th and 75th percentiles shown as shaded areas). The median spectrum of 100 biosonar calls is shown in D for comparison purposes. (E) Histogram of area under curve calculated from the spectra of FPVs and non-FPVs. The p-value of a Wilcoxon ranksum test and the *d* size-effect metric are provided. (F) histogram of peak frequency in FPVs and non-FPVs.

There were also differences between FPVs and non-FPVs regarding their harmonic-to-noise difference (HND, Fig. 4). The HND-metric is useful for quantifying spectral “smoothness”, and it is calculated as the difference between the observed spectrum and the same spectrum smoothened using a moving average filter (here a 5 point moving window applied to spectra calculated with 200-Hz frequency resolution; see Fig. 4A-D for illustration of the HND calculation in one FPV (Fig. 4A and C) and one non-FPV (Fig. 4B and D)). This method was originally proposed as a straightforward technique for studying “hoarseness” in human speech ^46^ and it has since been used in several studies on vocalizations produced by humans and other animal species (i.e. dog barks ^47^). Calculating the HND of FPVs and non-FPVs produced by *C. perspicillata* rendered statistical differences between the two syllable groups (Fig. 4E, p_ranksum=10_^-20^, median_FPVs_=0.45, median_non___FPVs_=0.39, small effect size (*d*= 0.3)) thus indicating that the spectra of syllables carrying fast periodicities is less smooth than that of non-FPVs.

**Figure 4.**
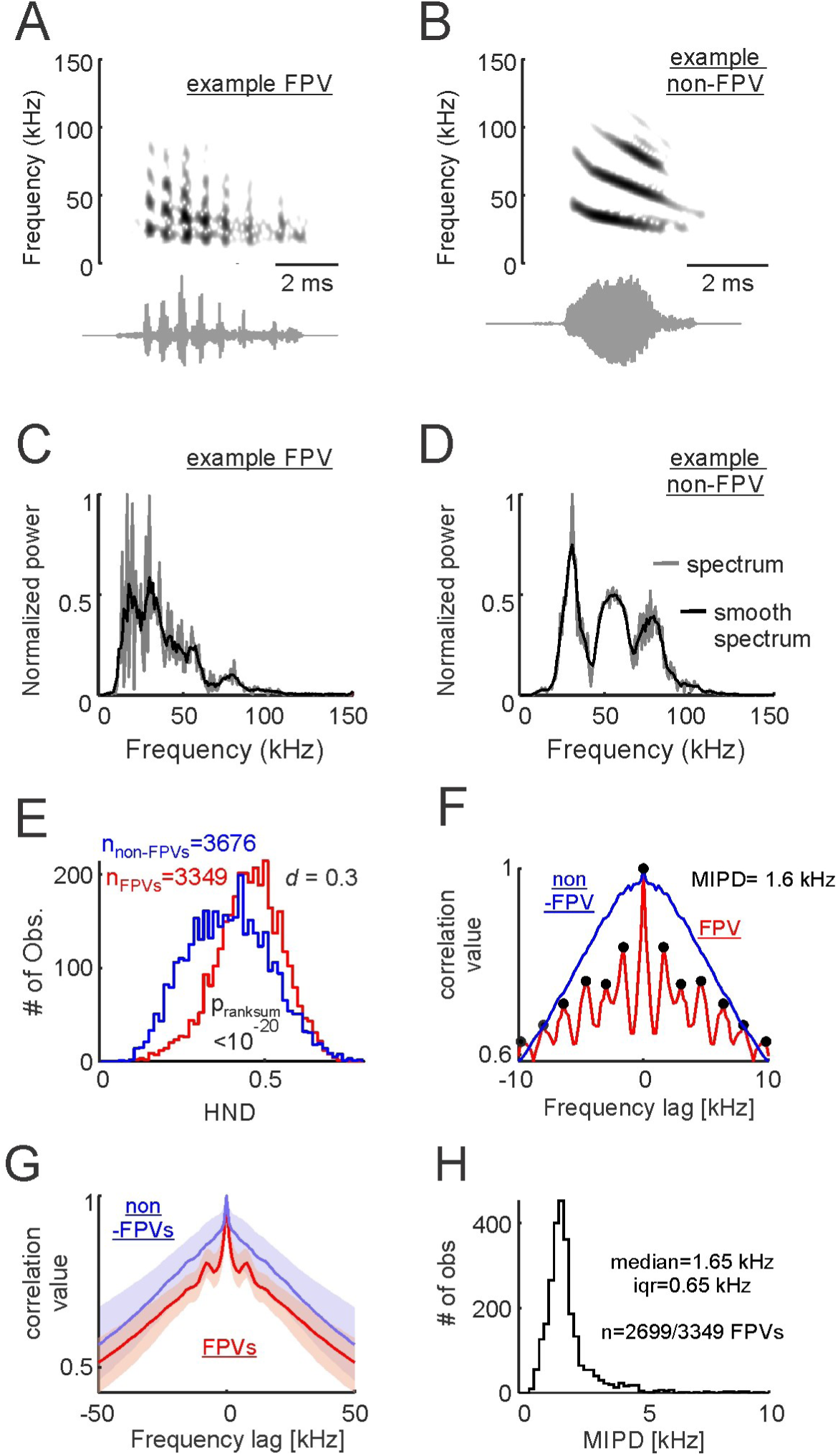
Spectral regularities are a characteristic feature of fast periodic vocalizations (FPVs). A and B show the spectrograms and waveforms of one example FPV and one non-FPV (A and B, respectively). C and D are the spectra of the same two example vocalizations. Note that the observed spectra (200-Hz frequency resolution) and “smooth” spectra are represented. Smooth spectra were calculated using a 5-point moving average. The difference between the observed and the smooth-spectra was used for harmonic-to-noise difference (HND) calculations. (E) Histograms of HND for FPVs and non-FPVs. The p-value of a Wilcoxon ranksum test and the d size-effect metric are provided. (F) Spectral autocorrelograms for the example FPV and non-FPV shown in C and D. Note that spectral regularities (i.e. peaks spaced at regular distances) occurred in the example FPV. In this example, the median inter-peak distance (MPID) was 1.6 kHz. (G) median spectral autocorrelograms for all FPVs and non-FPVs studied (25th and 75th percentiles shown as shaded areas). Note the side- peaks occurring in the population of FPVs. (H) Histogram of MPID for the FPVs in which more than one peak could be detected in the spectral autocorrelogram. Median and interquartile range are (iqr) given.

Note that the fact that FPVs had the least smooth spectra does not imply that their spectra were “irregular”. In fact, we observed that in the spectra of FPVs peaks occurred at regular intervals. The latter is illustrated in the spectrum autocorrelation function represented in figure 4F for one example FPV (same as in Fig. 4A, C). In this example autocorrelogram, local peaks (Matlab findpeaks function, peak prominence 0.025) could be detected every 1.6 kHz (i.e. 8 samples of the autocorrelogram of a 200-Hz resolution spectrum). Side-peaks indicating spectral regularity were also observed in the median autocorrelogram of all syllables labeled as FPVs by the SVM classifier, but not in the median autocorrelogram of syllables labeled as non-FPVs (Fig. 4G). The local-peak detection algorithm rendered more than 1 peak in 2699 out of 3349 syllables identified as FPVs (80.6%). In those syllables, we calculated the mean inter-peak distance (MIPD) from the autocorrelogram as a metric of spectral regularity. The MIPD distribution peaked at 1.6 kHz (Fig. 4H). Note that this value is close to best amplitude modulation frequency values determined by analyzing the temporal modulation spectrum (i.e. 1.75 kHz for all FPVs, see Fig. 3, but 1.65 kHz for the 2699 FPVs in which the MIPD could be measured). In fact, the results of a signrank test comparing the MIPD and best amplitude modulation frequencies of FPVs rendered no statistically significant differences (p_signrank_= 0.49). The latter suggests that temporal and spectral modulations are strongly linked to each other, a situation that is expected if one considers the spectral regularities observed as “sidebands” created by the presence of a modulating wave, or that the fast periodicities detected are simply a reflection of the signals’ fundamental frequency.

### 1.7 kHz: the missing fundamental of FPVs that does not trigger responses in the bat cochlea

Evidence presented thus far in this manuscript suggests that the 1.7 kHz periodicities measured in FPVs could be linked to the syllables’ fundamental frequency. We tested whether this fundamental frequency was missing or present in the syllables’ spectra. The latter was achieved by measuring the level (in dB SPL) at the FOIs (1.15 kHz-2.45 kHz). The level in this frequency range was obtained by computing the logarithm of the root-mean-square (RMS) of the filtered signals in the FOI range (3^rd^ order Butterworth filter), and by comparing the results with the RMS of a 94 dB SPL pure tone (1 kHz) produced by a calibrator (see methods). Overall, the level in the FOI range never exceeded 50 dB SPL, regardless of whether it was studied in FPVs or non-FPVs (Fig. 5A). The average FOI level for FPVs was −3.9 dB SPL while for non-FPVs the average level reached the 0.6 dB SPL. FOI level values were significantly higher in non- FPVs than in FPVs (p_ranksum_ < 10^-22^) although the size effect of this comparison indicated negligible effects (*d*=0.14). Note that the level values obtained in the FOI range were much lower than those obtained in the range from 20-21.3 kHz (mean level FPVs= 64.4 dB SPL, mean level non-FPVs= 61.2 dB SPL, p_ranksum <_ 10^-21^, *d*= 0.14). Overall, the low levels observed when syllables were filtered in the FOI range suggest that a “missing” fundamental frequency could be responsible for the temporal and spectral regularities measured in fast periodic distress syllables.

**Figure 5.**
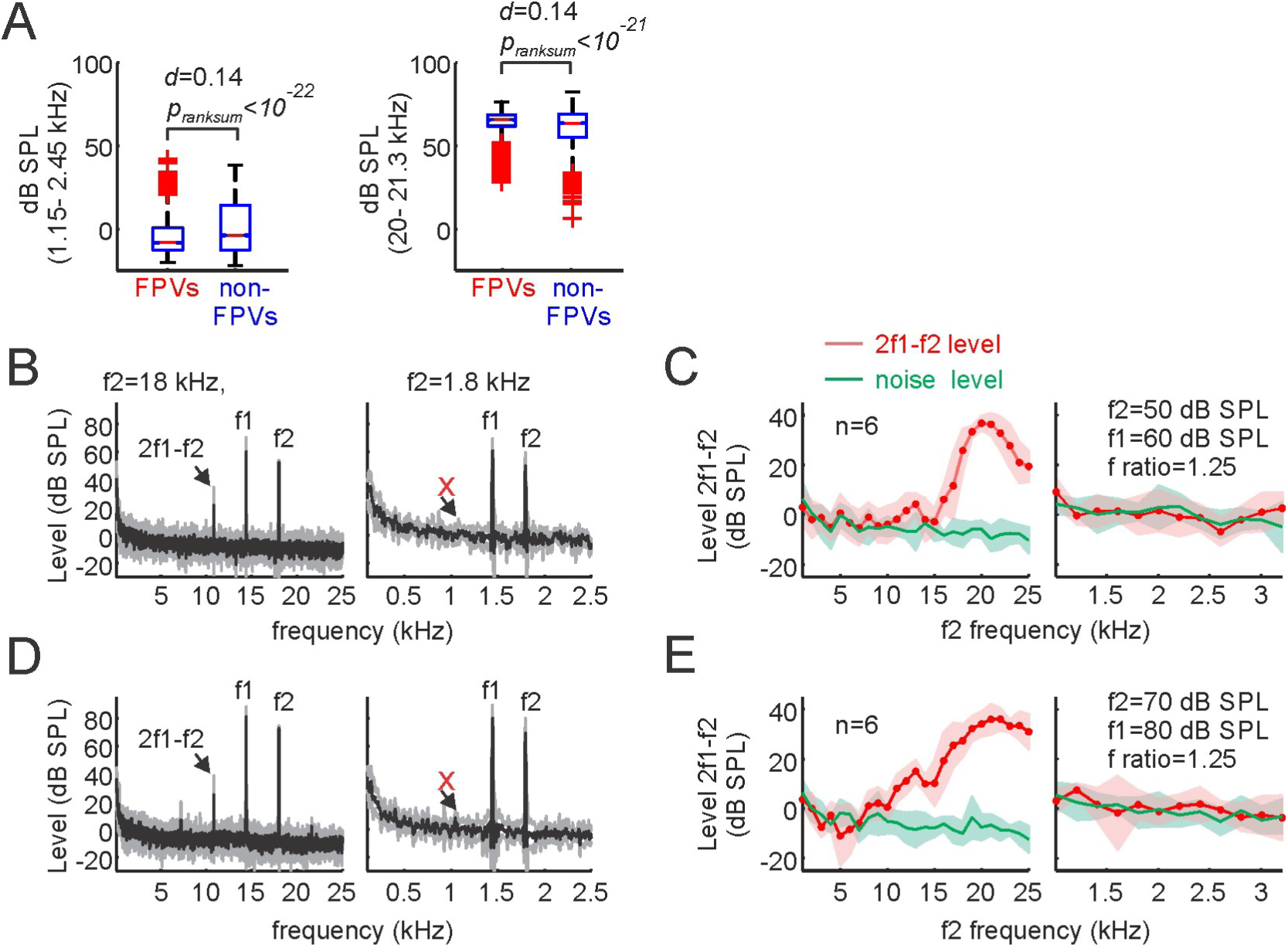
1.7 kHz is a missing fundamental that does not trigger responses in the bat cochlea. (A, left) Sound pressure level (in dB SPL) calculated in the range from 1.1 – 2.5 kHz. Note that this frequency range was not well represented in the sounds uttered, i.e. values were always below 50 dB SPL in FPVs and non-FPVs, and median SPLs were close to 0. The p-value of a Wilcoxon ranksum test and the d size-effect metric are provided. The same analysis is shown in the right panel of A for the frequency range between 21.1-22.5 kHz for comparison purposes. (B) average distortion product (DP) measurements obtained in six animals for f2 frequencies of 18 kHz and 1.8 kHz (left and right, respectively), when the f2 level was set to 50 dB SPL. Note that a strong cubic DP (2f2-f1) appeared when f2 was equal to18 kHz, but not when it was 1.8 kHz. (C) Coarse and high-resolution DPgrams obtained with frequency steps of 1 kHz and 200 Hz (left and right, respectively). Coarse DPgrams covered f2 frequencies from 1-25 kHz, while high- resolution DPgrams covered f2 frequencies from 1- 3.2 kHz. In high-resolution DPgrams, the DP level measured remained within the acoustic noise level, thus indicating poor hearing sensitivity for those frequencies. D and E show DP measurements similar to those depicted in B and C, but for stimulus levels of 80/70 dB SPL.

We also tested whether the bats’ ears were sensitive to frequencies ∼1.7 kHz (the putative fundamental frequency of FPVs). This was a necessary test, because we noticed that previous studies on *C. perspicillata*’s audiogram always measured hearing sensitivity at frequencies above 5 kHz ^48–50^. We measured the cochlear audiogram of 6 adult awake *C. perspicillata* (3 males, 3 females) by means of distortion product otoacoustic emissions (DPOAE). DPOAEs are a by- product of nonlinear ear mechanics and they represent a non-invasive objective method for measuring sensitivity and tuning of the cochlear amplifier ^51–53^. We focused on the cubic DPOAEs that occur at frequencies of 2f1-f2, were f2 and f1 represent the frequencies of two sounds produced simultaneously by loudspeakers placed closed to the bats’ tympanic membrane. The ratio between f1 and f2 was kept constant at 1.25 and there was a level difference of 10 dB between the sounds to optimize stimulus parameters (see Refs. ^53,54^). To test for the occurrence of DPOAEs, two stimulus level combinations were used (L1/L2 =50/60, 80/70 dB SPL). DPOAEs were measured with coarse and fine frequency steps, covering f2 frequencies between 1-25 kHz (steps of 1 kHz) and between 1-3.2 kHz (steps of 200 Hz), respectively.

As it can be seen in figure 5B and D, when f2/f1 sound pairs of 18/14.4 kHz were presented, a noticeable cubic distortion appeared, regardless of whether f2 was presented at 50 or 70 dB SPL (Fig. 5B and D (leftmost panels), respectively). However, no visible distortion product occurred in response to f2/f1 pairs of 1.8 and 1.44 kHz, regardless of the f2 level tested (Fig. 5B and D (rightmost panels)). Overall, high amplitude distortion products, indicating strong cochlear amplification, were visible only for f2 frequencies above 5 kHz (Fig. 5C and E, leftmost panels). In response to lower f2 frequencies, the distortion product amplitude fell within the acoustic noise level (Fig. 5C and E, rightmost panels). These results indicate that *C. perspicillata*’s cochlea is not well suited for dealing with faint low frequency sounds and can therefore not respond to potential 1.7 kHz fundamental frequencies of FPVs even if those frequencies were louder than 60 dB SPL, which is not the case according to our data (see SPL values in Fig. 5 A).

### The modulation power spectrum of distress syllables

We calculated the modulation power spectrum (MPS) of FPVs and non-FPVs (Fig. 6). The MPS is calculated from the 2D fast Fourier transform (FFT) of the syllables’ spectrogram (see below and methods). The MPS represents power in the two-dimensional space of temporal and spectral modulations and it has been used to study vocalizations in other highly vocal animal groups such as humans and birds ^55,56^.

**Figure 6.**
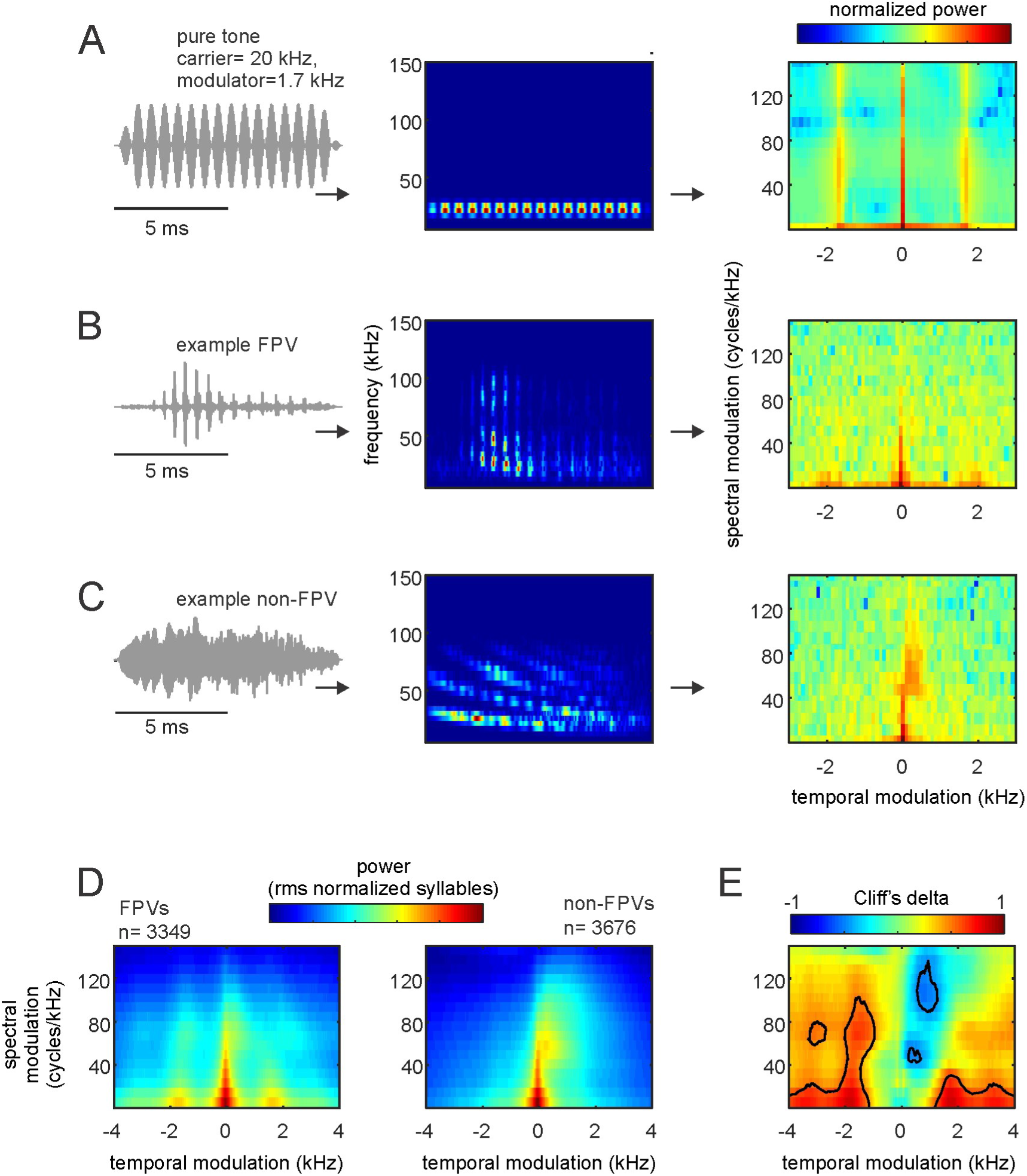
Modulation power spectra (MPS) of bat distress vocalizations. (A, from left to right) The waveform, spectrogram, and MPS of a pure tone temporally modulated at 1.7 kHz. Note that the temporal modulation appears in the MPS as side-peaks at −1.7 and 1.7 kHz. B and C show the waveform, spectrogram, and MPS for an example FPV and one non-FPV, respectively. Note that fast temporal modulations are visible in the modulation power spectrum of the example FPV. (D) Average MPS for the population of FPVs and non-FPVs studied. (E) Cliff’s delta (d) obtained after comparing the MPS of FPVs and non-FPVs. Contour lines indicate large size-effects (i.e. d>0.478), indicating consistent differences between the MPS of FPVs and non-FPVs.

We were interested in the MPS because of two reasons: (i) unlike classical acoustic analysis techniques (as those described in the preceding text), the MPS allows to quantify amplitude and spectral modulations simultaneously in each syllable ^56^; and (ii) filtering the MPS provides a robust technique for removing modulation components of the signal without changing other signal attributes. In our case, we were interested in determining whether the presence/absence of fast amplitude fluctuations had differential effects on the bats’ heart rate and neural responses (see below).

The oscillogram, spectrogram, and MPS of a 20 kHz pure tone modulated at 1.7 kHz, one example FPV, and one example non-FPV are shown in figure 6A-C, respectively. It can be noted that in both the amplitude modulated pure tone and the example FPV, more power occurred at temporal modulation frequencies close to 1.7 kHz. In the MPS, temporal modulations are represented in the positive and negative planes, with the former indicating the presence of downward frequency modulations and the latter corresponding to upward frequency modulations^55,56^. The presence of strong downward spectral modulations (hence positive values in the temporal modulation domain) is noticeable in the example non-FPV represented Figure 6C. This downward spectral modulation was strongest in the range between 50-80 kHz, corresponding to the call’s bandwidth.

As expected, pronounced power at temporal modulation frequencies close to 1.7 kHz was also evident when averaging MPS curves of all syllables classified as FPVs (n= 3349), but not in the average MPS of non-FPVs (n= 3676, Fig. 6D). We calculated the Cliff’s delta to assess the effect size of differences between the MPS of FPVs and non-FPVs (Fig. 6E). The comparison between the two syllable types was done for each temporal- and spectral-modulation combination in the MPSs. As mentioned in the preceding text, *d* values above 0.478 were considered as large effect size (contour lines in Fig. 6E) following previous studies ^45^. Overall, the values obtained from *d* calculations validated the existence of two main MPS differences between FPVs and non-FPVs: (i) faster temporal modulations in FPVs than in non-FPVs, and (ii) more pronounced downward spectral modulations in non-FPVs.

### Listening to fast periodic vocalizations boosts the bats’ heart rate

The acoustic analysis described in the previous sections revealed the presence of fast periodicities in 3349 out of 7025 syllables studied (47.7%). Such fast periodicities are likely due to a missing fundamental and the bats’ cochlea does not respond to the frequency of such missing fundamental in terms of nonlinear cochlear mechanics. To determine whether bats could actually perceive fast periodicities (or low pitch, see preceding text) we measured the heart rate (HR) response of awake animals while they listened to sequences of natural FPVs and their computer demodulated versions. Previous studies have shown that the bats’ HR increases when the animals are subject to fear conditioning, when they listen to aggression (versus non-aggression) calls, or after electric stimulation of the amygdala ^38,57,58^. We thus reasoned that heart rate could be a useful indicator of autonomic changes driven by the presence of fast periodicities in the sounds.

To determine whether fast periodicities had specific effects on the bats’ HR we filtered the MPS of three natural FPVs to produce their “demodulated” versions (Fig. 7). As stimuli, we chose one frequency modulated (FM) syllable (Fig. 7A), one syllable containing quasiconstant frequency and FM components (qCF-FM, Fig. 7B), and one containing sinusoidal frequency modulations (SFM, Fig. 7C). Note that in a previous study we reported that syllables containing FM components represent 94% of the distress vocalizations produced by *C. perspicillata*, SFMs (the next best represented group) amounted to ∼4% of the syllables analyzed, while qCF syllables represented less than 1% of the syllables studied ^35^.

**Figure 7.**
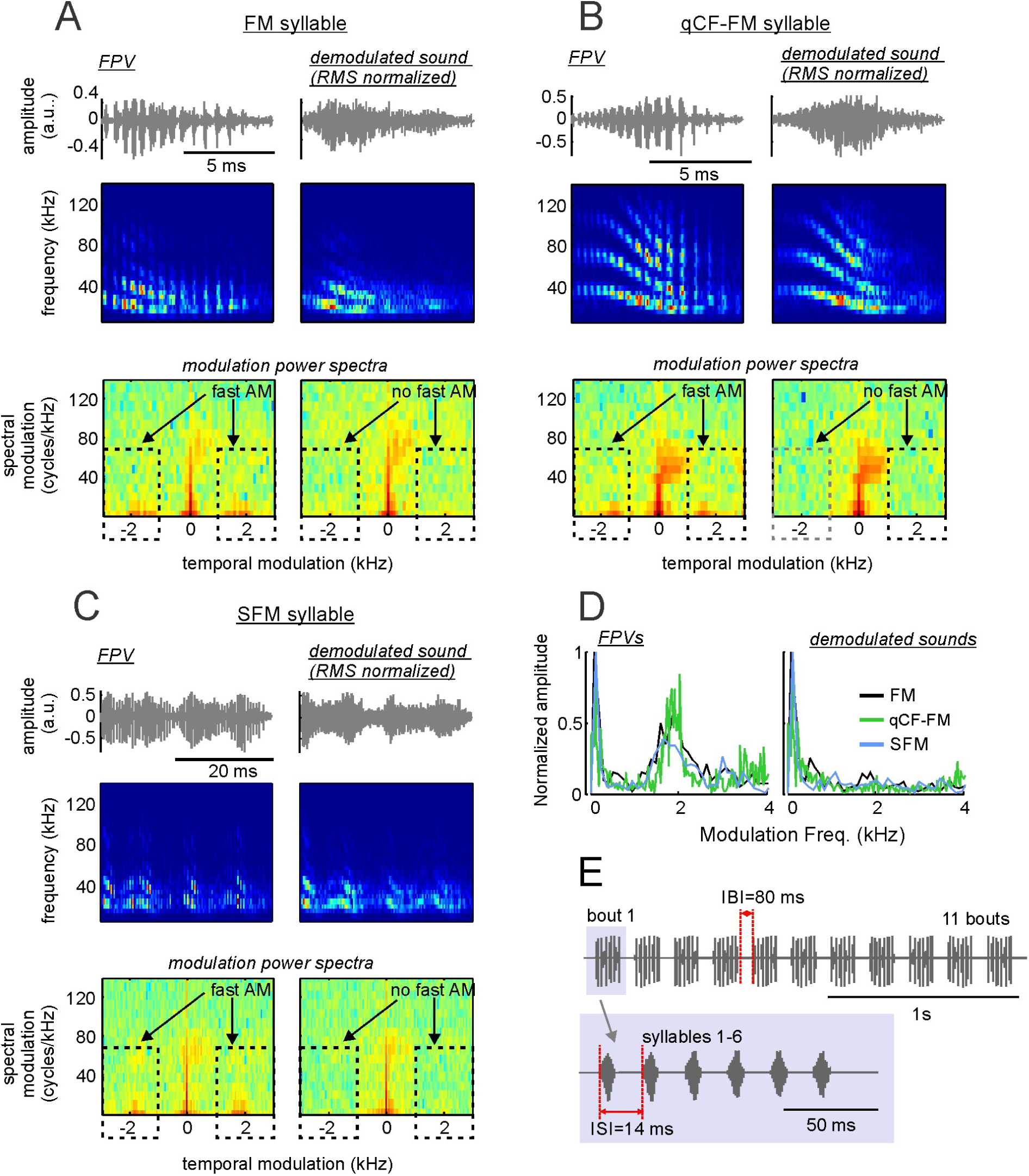
Removing fast periodicities by filtering modulation power spectra (MPS) of fast-periodic vocalizations (FPVs). A, left column shows the waveform, spectrogram, and MPS of a downward frequency modulated (FM) syllable containing fast periodicities. A, right column shows representations of the same sound after demodulation by filtering the MPS in the range marked with the dashed lines (see methods and supplementary figure S2). B and C show the same representations as A, but for one syllable containing quasiconstant-frequency and downward frequency modulated components (qCF-FM syllable, B) and one syllable composed of sinusoidal frequency modulations (SFM syllable, C). The three syllables represented in A-C and their demodulated treatments were used as stimuli for measuring electrocardiogram (ECG) and neural responses. (D) The temporal modulation spectra of the FM, qCF-FM, and SFM syllables (left) and their demodulated treatment (right). (E) Temporal arrangement of syllables in the sequences used as stimuli during ECG and neural measurement experiments. Altogether, six sequences were constructed, each composed of the same syllable repeated at inter-syllable intervals (ISIs) of 14 ms. The syllables formed 11 bouts that were repeated at inter-bout intervals (IBIs) of 80 ms.

The procedure used for MPS filtering was designed according to previous studies in humans ^56^ and is described in detail in the methods section and in the supplementary Figure S2, which shows the demodulation procedure for one of the sounds used as stimuli. Note that MPS filtering allows to modify certain features of the vocalizations without affecting others. In our case, we used MPS filtering for removing the syllables’ fast temporal periodicities without changing their spectro-temporal structure (see spectrograms in Fig. 7A-C). The result (Fig. 7A-D) was three pairs of natural FPVs and software-demodulated syllables (artificial non-FPVs) that were used as stimuli for measuring HR responses. Note that the TMS of artificial non-FPVs produced after MPS filtering resembles that of natural non-FPVs produced by the bats (Fig. 7D).

The final stimuli presented to the bats were sequences of either the natural FPVs or artificial-non FPVs in which the same sound was repeated 66 times in the form of 11 bouts (Fig. 7E, top panel), with 6 repetitions of the same syllable per bout (see Fig. 7E, bottom panel). The interbout interval was fixed to 80 ms and, within bouts, syllables were repeated at intervals of 14 ms. These parameters were chosen based on median values reported in a quantitative study on the structure of *C. perspicillata*’s distress sequences ^35^. HR changes in response to the acoustic signals described above were measured by attaching three electrodes (active, reference and ground) to the left and right sides of the chest and to the back of the animals, respectively (Fig. 8A). The resulting voltage differences were measured and the location of QRS complexes were detected automatically based on their amplitudes (Fig. 8B). Instantaneous HR was then calculated considering the interval between consecutive QRS complexes and expressed in beats/min. HR was measured from 5s before (baseline) until 10s after stimulus presentation.

**Figure 8.**
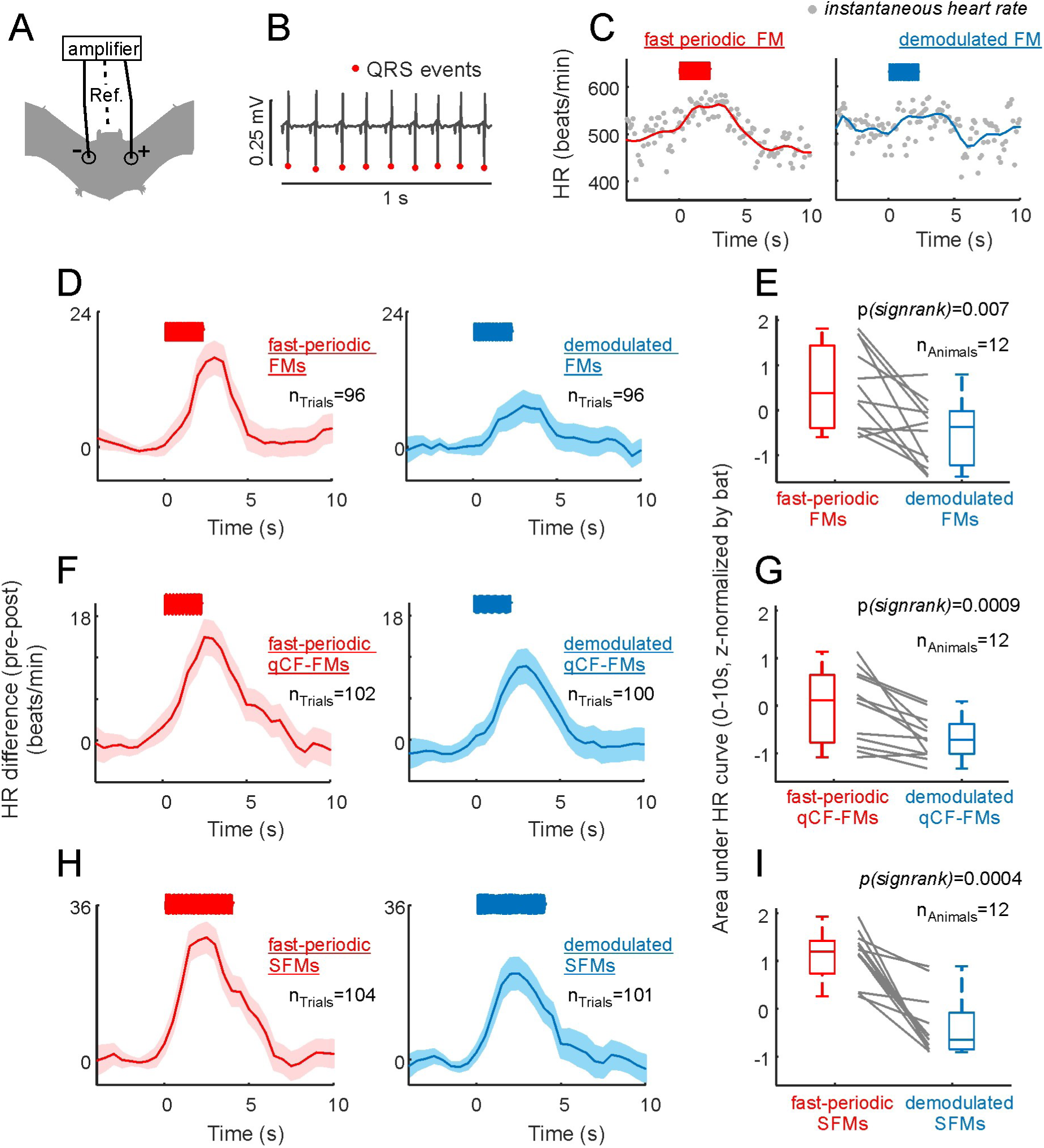
The presence of fast periodicities boosts the bats’ heart rate (HR). (A) Schematic representation of electrode positioning during electrocardiogram (ECG) measurements. (B) 1s- segment of ECG recording. The position of QRS events is indicated. (C) Instantaneous HR in two recording trials in which the fast-periodic FM syllable and its demodulated treatment were used as stimuli (left and right, respectively). HR-curves (solid-lines) were obtained by interpolation from the instantaneous HR data. (D) Average HR curves obtained considering all presentations of the fast-periodic FM syllable (left panel) and its demodulated treatment (right panel). Shaded areas indicate the standard error of the mean. Note that stronger HR responses followed the presentation of the fast-periodic FM. (E) Area under the average HR curve calculated in 12 awake bats in response to the two treatments of the FM syllable. Note that the fast-periodic treatment rendered the strongest HR responses. Area values have been z-scored for illustration purposes due to the HR variability across animals (z-scoring does not affect the results of paired statistics reported). F and G show similar results as those reported in D and E, but in response to the quasiconstant-frequency/ frequency-modulated (qCF-FM) syllable and its demodulated treatment. H and I show the results obtained when the sinusoidally frequency modulated syllable (SFM) was used as stimulus.

As mentioned in the preceding text, acoustic stimulation is known to increase the bats’ HR ^38,58^. This effect was also visible in our data, as illustrated in figure 8C (left) for two stimulation trials in which sequences of FM syllables were presented to an awake bat, with the syllables occurring either in their natural or demodulated forms (Fig. 8C left and right, respectively). Note that the example individual trials presented in Fig. 8C already point towards larger HR increments in response to fast-periodic than to demodulated sounds.

Average HR curves obtained by pooling data from all stimulation trials in all bats tested are shown in figure 8D, F& H (12 bats; 10 trials per animal and stimulus treatment, trials with movement artifacts were not considered, see methods). Regardless of the syllable analyzed, the natural treatment containing fast periodicities always produced higher HR increments than the demodulated treatment of the corresponding syllable. This was statistically validated by comparing the area under the HR curve in the first 10s after stimulation in each bat using paired statistics (Fig. 8E, G, I, n_bats_=12, FM syllable: p_signrank_ = 0.002, qCF-FM syllable: p_signrank_ = 0.00009, SFM syllable: p_signrank_ = 0.0004). Altogether the data obtained indicates that the presence of fast periodicities make signals more effective in boosting the bats’ HR. The latter points towards a role of fast amplitude fluctuations for signaling distress.

### Listening to fast periodic vocalizations triggers frequency-following responses in the bat brain

Our results show clear evidence on the existence of fast periodicities in bat distress vocalizations. Such fast periodicities accelerate the bats’ heart rate. For the latter to occur, fast periodicities have to be encoded in the bats’ brain. We investigated whether frequency-following responses (FFRs) occurred in response to FPVs. FFRs appear as rhythmic brain signals occurring at the same frequency of the sensory input (i.e. 1.7 kHz in FPVs). In humans and other animal species, FFRs have been used to study the auditory system’s ability to process temporal periodicities ^59–63^ and it is known that FFRs can occur in response to missing fundamental frequencies ^64^.

FFRs were studied by measuring intracortical electroencephalogram signals (iEEG, active electrode placed over the auditory cortex) in 11 head-restrained, awake bats. As stimuli, the same sequences of natural FPVs and demodulated syllables used for measuring HR (see Fig. 7) were presented. Figure 9 A-B show the average iEEG obtained across animals in response to the sequence of fast-periodic FMs (Fig. 9A, top panel) and demodulated FMs (Fig. 9B, top panel). Spectrograms of the signals recorded were calculated to assess the power relations at the FOIs in the neural responses to fast-periodic FMs and demodulated FMs (Fig. 9 A-B, bottom panels). From the spectrografic representations, it is clear that FPV-evoked responses had more power in frequencies close to 1.7 kHz. The latter becomes obvious after subtracting the two spectrograms (Fig. 9C). Such pronounced power at frequencies close to 1.7 kHz is likely related to the occurrence of an FFR that represents the bat auditory system’s ability to extract fast periodicities. The FFR can also be visualized as fast fluctuations in the neural signals obtained after averaging the 20 ms time-window following the presentation of each syllable across sequences, trials and animals (Fig. 9D, n=24950 responses to the FM syllable).

**Figure 9.**
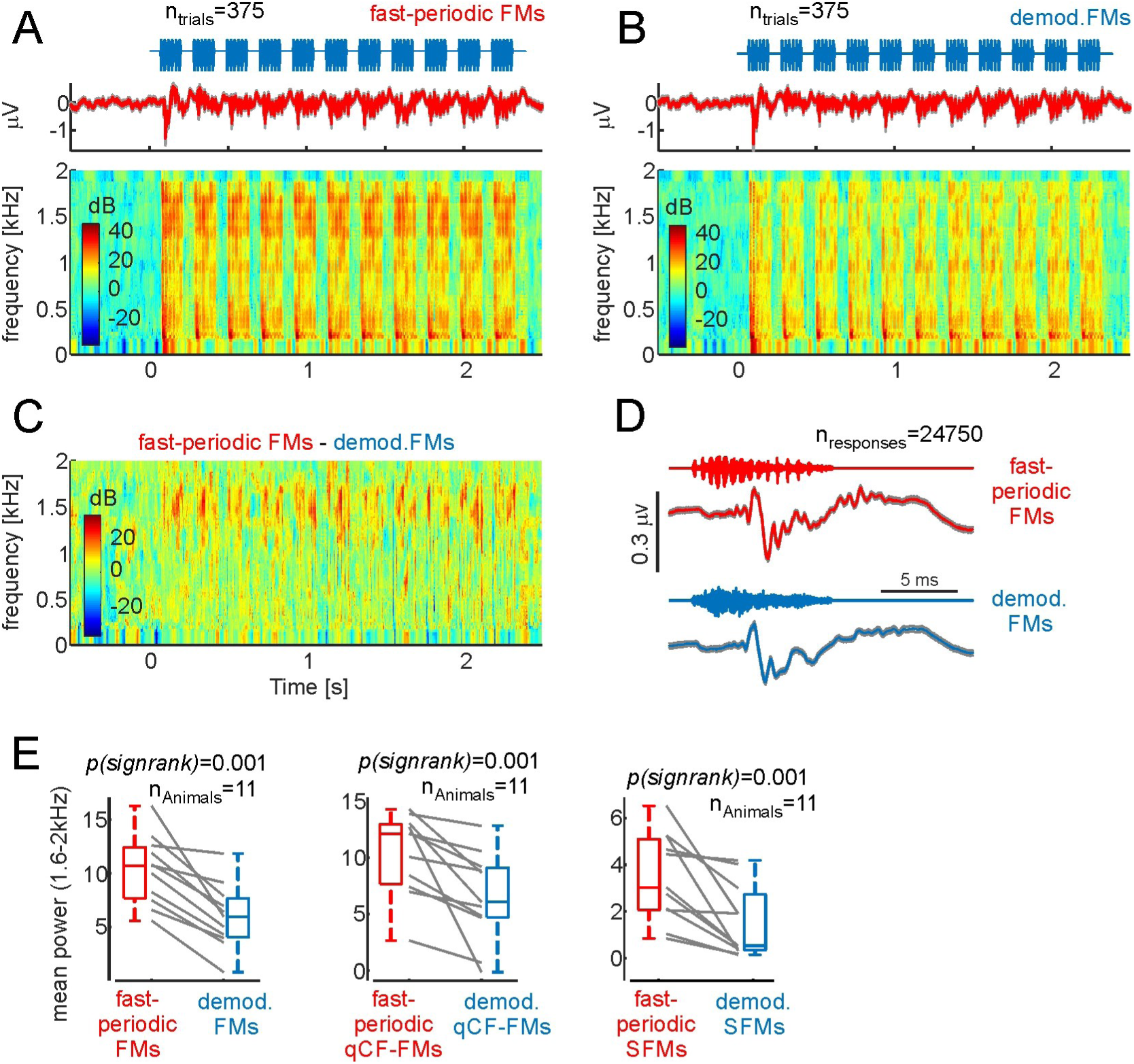
Fast periodic vocalizations trigger frequency following responses in the bat brain. (A) Average neural responses to a sequence of fast periodic FM syllables across all trials and animals (n=11) studied. Responses are represented as voltage vs. time and in the form of neural spectrograms. (B) Same as A but in response to the sequence of demodulated FMs. Note that at frequencies ∼ 1.7 kHz, more power occurred in response to fast-periodic than to demodulated FMs. (C) Difference between the neural spectrograms depicted in A and B. (D) Voltage fluctuations obtained after averaging neural responses to each fast-periodic FM (red) and each demodulated FM syllable (blue) across trials and animals. Note that responses to fast periodic FMs carried faster modulations than those obtained in response to demodulated FMs. (E) Mean power in the range from 1.6-2 kHz across animals and sequences studied. In each animal fast periodic syllables rendered higher power than their demodulated treatment.

To statistically validate the presence of an FFR, the average power (across time) at the FOIs was calculated in the neural spectrograms obtained for each animal in response to each of the six sequences studied (sequences composed of: (1) modulated FM syllable, (2) demodulated FM syllable, (3) modulated qCF-FM syllable, (4) demodulated qCF-FM syllable, (5) modulated SFM syllable, (6) demodulated SFM syllable). Average neural spectrograms pertaining responses to the qCF-FM and SFM syllables can be found supplementary figure S3. At the population level, when considering the power of the neural responses in the range from 1.6 to 2 kHz, there were significant differences between responses to natural FPVs and demodulated FPVs in all three cases studied (p_signrank_ = 0.001 for all three cases studied, Fig. 9E). The latter indicates that the bat auditory system is capable of representing the fast periodicities found in FPVs using a temporal code.

## Discussion

The main aim of this article was to study the temporal modulation pattern of distress syllables produced by bats, a highly vocal animal group. We tested the idea that fast amplitude fluctuations could be a generalized trait of mammalian vocalizations for signaling distress. If this thesis were true, then, a large percentage of bat distress vocalizations should carry fast periodicities.

Four main observations support our driving hypothesis. (i) Almost half (47.7%) of the distress syllables produced by bats (species *C. perspillata*) carry superfast periodicities at ∼ 1.75 kHz. (ii) These fast temporal periodicities could be linked to a missing fundamental frequency that does not evoke a response in the bats’ cochlea based on non-linear mechanics. (iii) Vocalizations carrying fast periodicities produce larger heart rate increments than their demodulated versions, thus suggesting that fast periodicities can indeed be used for conveying alarmfullness in bats. (iv) Fast periodic vocalizations evoke frequency following responses, indicating that the bats’ auditory system can represent fast amplitude modulations.

### Comparison with previous studies

Several studies in bats and other animal species have characterized the amplitude modulation pattern of natural vocalizations. A recent study in the bat species Phyllostomus discolor (a sister species of *C. perspicillata*) described vocalizations carrying amplitude modulations at rates close to 130 Hz ^65^. Periodicity values below 500 Hz also have been described in previous studies in frogs and birds ^66–71^ . In humans, amplitude fluctuations occur in screamed vocalizations in the form of roughness at temporal modulation at values close to 100 Hz ^1^. The periodicity values reported in the present article reach 1.7 kHz, that is, 17 times faster than modulation rates reported in human screams and at least 8 times faster than modulation rates reported in other vertebrates ^67,70^, including other bat species ^65^.

Bats and humans are phylogenetically distant species that do not share common ecological niches. Yet, in both species fast temporal periodicities are present in fearful vocalizations. Note that the roughness reported in human screams was obtained by comparing screamed and non- screamed utterances after pitch-normalizing the latter ^1^. However, a recent study showed that in humans, identifying sounds as screams is also linked to high pitch values (pitch in screams is higher than in the neutral sounds ^72^). In bats, fast periodicities appear to be linked to a missing fundamental that could be interpreted as an acoustic correlate of “pitch”. In this sense, bats differ from humans in the strategy they use for achieving high periodicities in the sounds emitted. While the latter increase their pitch and temporally modulate their sounds when screaming, the former simply rely on low pitched sounds to make vocalizations more arousing, as assessed here with HR measurements.

Although pitch is a likely candidate for explaining the fast periodicities observed in our dataset, it is worth mentioning that fast temporally periodic structures leading to complex spectra could also be related to non-linear phonation phenomena such as “torus” and “deterministic chaos” ^73–75^. In bats, these two explanations (pitch and non-linear phenomena) might not be mutually exclusive. Pitch is a purely perceptual measure that can be related to sounds’ fundamental frequency, while non-linear phenomena are identified based on the sounds’ entire spectrograms. Previous studies in bats have reported the occurrence of non-linear phenomena ^76,77^. Particularly deterministic chaos appears to be linked specifically to high aggression contexts ^76^.

The occurrence of non-linearities during vocalization production is ubiquitous in vertebrates ^75–80^. It has been argued that non-linear sounds result from saturation in the vocal production apparatus, and that their generation does not require complex neural control mechanisms ^47,73,75^. Yet, non-linearities are exploited by several vertebrate species, including human infants, to capture the listeners’ attention, due to the non-predictable structure of the sounds uttered (for review see Ref. ^73^), and to prevent behavioral habituation ^81^. Bats could profit from the presence of spectro-temporally complex sounds (like the FPVs reported here) within distress broadcasts. Most bat distress sequences are long (> 1s) and repetitive, since the same syllable spectro- temporal design is used throughout the broadcast ^35,82^. One could speculate that the occurrence of fast-periodic syllables could re-capture the listeners’ attention thus preventing rapid neuronal adaptation of responses to the individual syllables.

### Possible neural mechanisms for fast periodicity extraction

Our data shows that the presence of fast periodicities accelerates the heart rate of awake bats. This indicates that fast periodicities are extracted somehow in the bats’ brain. Note that the audiogram of most bat species is shifted towards high (ultrasonic, >20 kHz) frequencies. For example, auditory thresholds in *C. perspicillata* have values above 70 dB SPL for frequencies below 10 kHz ^48,50^. In fact, the DPOAE measurements presented here showed no cochlear responses to low frequency sounds below 5 kHz. According to our data, the sound pressure level of the missing fundamental is very low (average values ∼ 0 dB SPL), which further hampers its representation at the cochlear level.

Two possibilities come to mind when thinking about neural strategies for coding fast periodicities: (i) the use of spectral harmonic codes and (ii) temporal codes ^83,84^. Spectral harmonic coding does not depend on a region of the cochlea being able to extract low fundamental frequencies but rather on the ability of the cochlea to resolve closely placed harmonics of that fundamental ^67^. Whether the cochlea of *C. perspicillata* can resolve harmonics separated by 1.7 kHz remains to be tested. It has been argued that the exact periodicity value at which a switch from temporal to spectral coding occurs might differ across species ^67^. At least in the auditory nerve of squirrel monkeys, spiking activity can statistically lock to the occurrence of periodicity cycles for frequencies up to 5 kHz (temporal coding ^85^). If the same is assumed for bats, then the 1.7 kHz modulation shown here could be encoded in auditory nerve activity patterns. The FFR measurements reported in this manuscript are in agreement with this idea.

We show that surface potentials represent the fast periodicities occurring at frequencies ∼ 1700 kHz based on a temporal code. Note that FFRs in response to amplitude fluctuations faster than 1 kHz are not unique to bats ^60^. Our recordings were based on surface potentials (iEEG) that are suited for studying whole-brain activity, but are not ideal for identification of possible generators contributing to the measured signal. Previous studies measuring FFRs in humans concluded that FFRs obtained in response to fast frequencies (i.e. >100 Hz) typically result from activity in subcortical structures ^62,86,87^. The same could be true for *C. perspicillata*, since in this species most auditory cortex neurons cannot track amplitude modulations above 20 Hz ^41,88^, even though field potentials measured at the cortical level do entrain to faster acoustic rhythms ^39,40^.

In *C. perspicillata*’s AC, there are also neurons that represent amplitude fluctuations by means of rate-coding. Rate coding involves an increased spike count in response to certain modulation rates and it has been suggested as a possible candidate for pitch representation in the brain ^89,90^. With our current data we cannot assess whether specialized rate-coders exist in C. perspicillta’s AC, which could play a role in the representation of superfast periodicities. Future studies measuring spiking activity are needed for the latter. Note that structures outside the classical ascending auditory pathway could also be involved in the representation of fast periodic vocalizations. For example, the amygdala is a likely candidate for providing such representations. In humans, this structure is differentially activated by screamed and non- screamed sounds ^1^. In bats, electric stimulation of the amygdala triggers changes in heart rate ^57^. It is thus plausible to suggest an involvement of the amygdala in the elevated HRs reported in this article in response to FPVs.

Taken together, the findings reported in this manuscript indicate that ultrasonic animals (such as the bat species *C. perspicillata*) can exploit the low frequency portion of the soundscape to add fast temporal periodicities and frequency harmonics to the sounds produced. Such periodicities are represented by the bats’ auditory system as a frequency following response and they accelerate the bats’ heart rate, an autonomic response to alarmfullness that could be instrumental for the bats’ survival.

## Methods

### Distress call recording and analysis

All the experiments described in this article were carried out in accordance with current laws for animal experimentations in Germany (Regierungspräsidium Darmstadt, permit # F104/57) and with the declaration of Helsinki. Distress vocalizations were recorded from 13 adult bats (6 females and 7 males) of the species *C. perspicillata*. Bats were captured in a breeding colony at the Institute for Cell Biology and Neuroscience (Frankfurt University) and brought one by one into an acoustically isolated chamber where the distress vocalization recordings took place. Methods used in this article for recording distress calls have been described elsewhere ^35^. In a previous article, we focused in studying the properties of distress “sequences” without considering the presence of fast periodic vocalizations within individual syllables. The latter is the main focus of this paper.

For acoustic recordings animals were hand-held with their face pointing straight into a microphone (Brüel&Kjaer, ¼-inch Microphone 4135, Microphone Preamplifier 2670) located at 1.5 m from the bat. To encourage the production of distress calls, the researcher holding the animal softly caressed the neck-skin of the bats. Recordings lasted up to 3 min per bat. The recording microphone was powered via a custom-built microphone amplifier and connected to a commercially available sound acquisition system (UltraSoundGate 116Hm mobile recording interface, + Recorder Software, Avisoft Bioacoustics, Germany) for sound digitization at 300 kHz (16-bit precision). Digitized signals were stored in a computer for offline analysis using the Avisoft SAS Lab Pro software (v.5.2 Avisoft Bioacoustics, Germany). The temporal position of individual “syllables” in each recording was automatically detected using an amplitude threshold of 4.1% of the maximum recording amplitude allowed when recording with the microphone amplifier gain set to the minimum. A syllable was defined as a fluctuation in amplitude in which the signal level did not drop below the amplitude threshold criterion (the 4.1% mentioned above) for a period of at least 1ms. Amplitude detection was manually revised for each syllable to ensure the accuracy of the results.

The temporal modulation spectrum (TMS), frequency spectrum, spectrogram, and modulation power spectrum (MPS) of each syllable were calculated and used for acoustic analysis. TMS was calculated as the FFT of each syllable’s amplitude envelope (secant method, temporal resolution = 0.1 ms). Frequency spectra were calculated as the FFT of each syllable’s waveform and interpolated to a resolution of 200 Hz for averaging purposes, using a linear interpolant. Short time Fourier transforms (STFTs) were calculated on zero-padded signals (0.5 s padding) using the following parameters: window length = 64, number of FFT points = 64, hop =1. The sampling rate was equal to 300 kHz. Zero-padding was necessary for obtaining STFTs of similar temporal and spectral resolutions across the syllables studied. The STFTs obtained were then used for computing modulation power spectra (see below).

For syllable classification based on their TMS, a binary support vector machine (SVM) classifier was used. The SVM classifier was trained (*fitcsvm* function, rbf kernel, Matlab 2018, no standardization) using the TMS of 100 vocalizations: 50 vocalizations contained pronounced periodicities in the range from 1.1-2.5 kHz, and another 50 vocalizations had no pronounced power in their TMS for that frequency range (see training TMS sets in supplementary figure S2). The vocalizations chosen for the training sets were randomly picked after visual inspection of the entire dataset. The model cross-validation error (calculated using 10-fold cross-validation) amounted to 2%.

Harmonic to noise differences (HND) were used to complement classic spectral analysis. HNDs were calculated as the difference between the observed- and smooth-FFT of each syllable. The smooth-FFT was obtained using a 5-point moving average filter that removed peaks in the observed-FFTs. All FFTs had a frequency resolution of 200 Hz. The latter was achieved by linear interpolation of the FFTs obtained from each sound. This was a necessary step, since frequency resolution is linked to sound length. The HND of each sound was equal to the maximum absolute difference between the observed and smoothed FFTs. To characterize the presence of spectral regularities spectral autocorrelograms were used. To that end, the spectrum of each syllable was autocorrelated for frequency of up to ±10 kHz. The median interpeak distance (MIPD) was used to measure regularity values. MIPDs were obtained after detecting local peaks in the autocorrelograms’ local maxima using the peakseek function (peak prominence = 0.025). This procedure was effective (i.e. it detected more than one peak) in 2699 out of 3349 FPVs detected (80.6%). In the remaining FPVs, the spectra were too noisy for local peak detection.

### Computing modulation power spectra

The MPS represents each syllable in the spectral and temporal modulation domains (see Fig. 6) and it was calculated as the 2D-FFT of the log-transformed STFTs. The absolute value of the 2D-FFT was then squared and log-transformed to produce the MPS. Note that spectrogram parameters (i.e. number of FFT points, window length and hop, see above) were chosen so that the temporal resolution of spectrographic representations was precise enough for representing periodicity values around 1.7 kHz in the temporal modulation domain. Using a larger window size could have resulted in periodicity representations in the spectral modulation domain, rather than in the temporal domain. STFTs were calculated using a linear frequency axis thus rendering spectral modulations in the MPS expressed in cycles/kHz. STFTs obtained with a log frequency axis result in spectral modulations given in cycles/octave. Previous studies have suggested that MPS representations in cycles/kHz are useful when dealing with harmonic sounds, as it was the case here ^56^.

For constructing the stimuli used in ECG and iEEG experiments, three natural fast periodic syllables (see Fig. 7 and Fig. S2) were demodulated using an MPS filtering algorithm similar to that described in previous studies in humans ^56^. Before MPS filtering the three syllables used as stimuli were downsampled to 192 kHz. MPS filtering was achieved by nullifying all MPS-power across spectral modulations in the temporal modulation range from −1 to −4 kHz and from 1 to 4 kHz. This temporal modulation range covered the fast periodicities of interest, occurring at frequencies at ∼1.7 kHz. The filtered MPS was then exponentiated, root-mean squared, and transformed into a matrix of complex numbers, built taking into account the phase matrix obtained from the 2D-FFT of the original sounds. The resulting matrix was then transformed into an STFT using an inverse FFT2 procedure. The resulting STFT was then exponentiated and transformed into a sound waveform using an inverse STFT, implemented based on an inverse FFT and the weighted-overlap-add method ^91^. The new demodulated sound and the natural FPV from which it derived were then root-mean-square normalized to avoid level differences. The sound synthesis procedures described above involve inverse STFTs that could be affected by time-frequency trade-offs. To quantify possible errors during sound synthesis using inverse STFTs we used a method proposed in previous studies ^56^, in which the difference between the desired and observed STFTs are squared and divided by the desired STFT. The observed STFT is obtained as the STFT of the newly synthetized sound, while the desired STFT is obtained after exponentiation of the outcome of the inverse STFT obtained from the filtered MPS. For all three sounds used, the synthesis error was below 2%.

### Setup for ECG measurements

The natural FPVs and their demodulated versions obtained from MPS filtering were used to build acoustic stimulation sequences as described in figure 7. For sequence building, the start and end of each syllable was multiplied by a linear fading window of 0.2 ms to avoid acoustic artefacts during stimulation. Sounds were synthetized in MATLAB 2015 (The MathWorks, Inc., Natick, Massachusetts, United States), produced through a sound card (RME Fireface 400, sampling rate = 192 kHz), amplified (Rotel power amplifier, RB-850) and played from a speaker (NeoCD 1.0 Ribbon Tweeter; Fuontek Electronics, China) placed 15 cm in front of the bats’ nose. The RMS level of the 6 syllables (3 natural FPVs and 3 demodulated syllables) when produced by the speaker spanned between 68.6 and 70.5 dB SPL (mean = 69.3 dB SPL, std= 0.7 dB SPL). To prevent adaptation, sequences were played randomly at intervals of 3 min between each sequence presentation. All measurements were conducted inside a soundproofed chamber.

Electrocardiogram (ECG) measurements were conducted in 12 awake animals (5 females, 7 males) placed on a custom-built holder similar to those used in electrophysiology experiments ^39,92,93^. ECG signals were obtained by placing three electrodes (active, reference, and ground) on the bats’ chest and back (see Fig. 8A). We found this configuration to be more stable for ECG recordings in awake bats than configurations involving the thumbs and legs ^94^. The three electrodes were attached to the inside of a custom-built Velcro belt. Electrolytic gel (Supervisc, EasyCap GMBH, Germany) was used to improve the contact between electrodes and skin. After the experiments, the skin was carefully cleaned using cotton-swabs and water.

Measuring electrodes were attached to a pre-amplifier/amplifier system (EX1 Differential amplifier, Dagan Corporation). Signals were amplified (gain=50) and band-pass filtered by the recording amplifier between 0.1 kHz and 1 kHz. ECG signals were digitized using the same sound card used for acoustic stimulation (see above), down-sampled to 9.6 kHz, and stored in a computer for offline analysis. To facilitate the automatic detection of QRS complexes (see below) the signal was adjusted so that the largest amplitude deflection recorded had a negative sign. The instantaneous heart rate was calculated as the inverse of the interval between consecutive QRS complexes multiplied by 60, to express it in beats/min. QRS complexes were identified by setting an amplitude threshold that detected the QRS events as “spikes” whose amplitude was larger than at least two standard deviations of the noise level calculated from the envelope of the ECG signal.

Altogether, we presented 10 trials of each syllable and stimulus treatment (natural and demodulated) amounting to six different conditions. The 10 trials were split into two blocks with a break of 10 min between blocks during which water was offered to the animals. Awake bats occasionally moved during the recordings. Trials that contained movement artifacts were excluded from the analysis. In ECG recordings, movement artifacts appear as signal peaks (spikes) occurring at intervals shorter than 62 ms thus producing instantaneous frequencies above 960 beats/min. This value had been used in a previous article for movement detection ^94^ and was used here for trial rejection. Overall, trials contaminated with movement artifacts represented 17% for the total number of trials gathered across all animals and stimuli tested (599/720). To average HR measurements across trials, instantaneous HR values were linearly interpolated with a temporal resolution 0.5 s.

### Setup for DPOAE measurements

DPOAEs were recorded in 6 adult awake *C. perspicillata* (3 males, 3 females) in a soundproofed chamber. To ensure that bats were not able to move during the recordings, their heads were fixed by holding a metal rod attached to the scalp. The surgical procedure for metal rod fixation has been described elsewhere ^41,92,95^. Briefly, in fully anesthetized bats (Ketamine (10 mg *kg^−1^ Ketavet, Pfizer) and Xylazine (38 mg *kg^−1^ Rompun, Bayer)), the skin and muscles covering the scalp were removed. The scalp surface was cleaned, and a custom-made metal rod (1 cm length, 0.1 cm diameter) was then glued to the skull using dental cement (Paladur, Heraeus Kulzer GmbH).

The DPOAE setup followed the specifications described in previous studies ^52,53^. To measure DPOAEs, an acoustic coupler was placed in the outer ear canal at a distance of about 0.3-1.0 mm from the tympanum under visual control (Zeiss OPMI 1-FR binocular, Carl Zeiss AG, Jena, Germany). The coupler consisted of three acoustic channels that converged at the coupler’s tip. Two of the coupler channels were connected to reversely driven condenser microphones used as loudspeakers (1/2“, MTG MK202, Microtech Gefell GmbH, Gefell, Germany) and the third channel contained a sensitive microphone (1/4“, B&K 4939, Brüel & Kjær, Nærum, Denmark) for recording DPOAEs. A soundcard was used to generate the two pure tone stimuli and to record DPOAEs (RME fireface UC, RME Audio AG, Haimhausen, Germany; sampling rate: 192 kHz). Data acquisition and data analysis programs were written in MATLAB (MATLAB 2015b, MathWorks Inc.). The sound system was calibrated in situ before each measurement using white noise. DPOAEs were recorded by varying the stimulus frequency f2 between 1 and 25 kHz (1 kHz steps) and between 1 and 3.2 kHz (0.2 kHz steps) to obtain DPOAE data at coarse and fine frequency resolution, respectively. The ratio between f2 and f1 frequencies was kept constant at 1.25. Two f2 levels were tested: 50 and 70dB SPL (f1 level= 60 and 80 dB SPL, respectively). To calculate DPOAE amplitudes, FFT-analysis was performed from 100 averages of the time signal acquired based on 8192-point epochs. The noise floor was calculated as the arithmetic mean of the amplitude of 20 points in the spectrum taken on either side of the DPOAE frequency within a 100 Hz frequency span. This method yielded DPgrams (plots of DPOAE amplitude versus f2 frequency) that were used to confirm that hearing deteriorates in *C. perspicillata* for frequencies below 5 kHz.

### Setup for neurophysiology measurements

Intracranial electroencephalogram (iEEG) signals were measured to assess the occurrence of frequency following responses that could represent the fast periodicities found in FPVs. iEEGs were measured in fully awake, head-restrained animals. The head of the bats was immobilized by holding a metal rod attached to the scalp. The surgical procedures used to attach the metal rod were similar to those described in the preceding text (see methods for DPOAE measurements). iEEGs were obtained using silver wires placed below the scalp. Three wires were used (active reference and ground). In each animal, the active electrode was placed over the primary auditory cortex, the reference was placed on a similar rostro-ventral position as the active electrode but close to the midline and the ground electrode was placed over the cerebellum. The location of the primary auditory cortex was estimated using external landmarks such as the medial cerebral artery and the pseudo-central sulcus ^92,93,96^.

Measuring electrodes were attached to a pre-amplifier/amplifier system (EX1 Differential amplifier, Dagan Corporation). Signals were amplified (gain=50) and band-pass filtered by the recording amplifier between 0.1 kHz and 5 kHz. iEEG signals were digitized using the same sound card used for acoustic stimulation (see above), downsampled to 9.6 kHz, and stored in a computer for offline analysis. The multi-taper method was used to estimate spectral power in the iEEG signals recorded ^97^ (5 tapers, time-bandwidth product of 3). Neural signals are known to follow a power rule by which high frequencies contain less power than lower frequencies. For better visualization of the power at high frequencies (i.e. 1.7 kHz), spectrograms were corrected by subtracting the average power at each frequency during time periods of 1.5 s, in which no acoustic stimulation was presented.

