## supplementary figures S1-S3 for "Superfast periodicities in distress vocalizations emitted by bats"

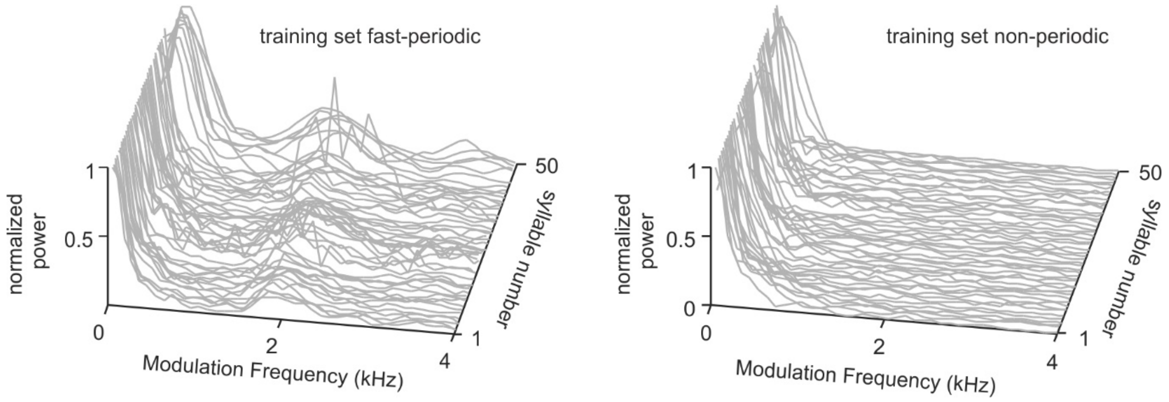


Figure S1. Temporal modulation spectra of vocalizations used to train the support vector machine classifier. The fast-periodic set includes 50 randomly chosen vocalizations containing fast periodicities at ~ 1.5 - 2 kHz. The non-periodic set includes 50 randomly chosen vocalizations that contained low power in the 1.5-2 kHz range.

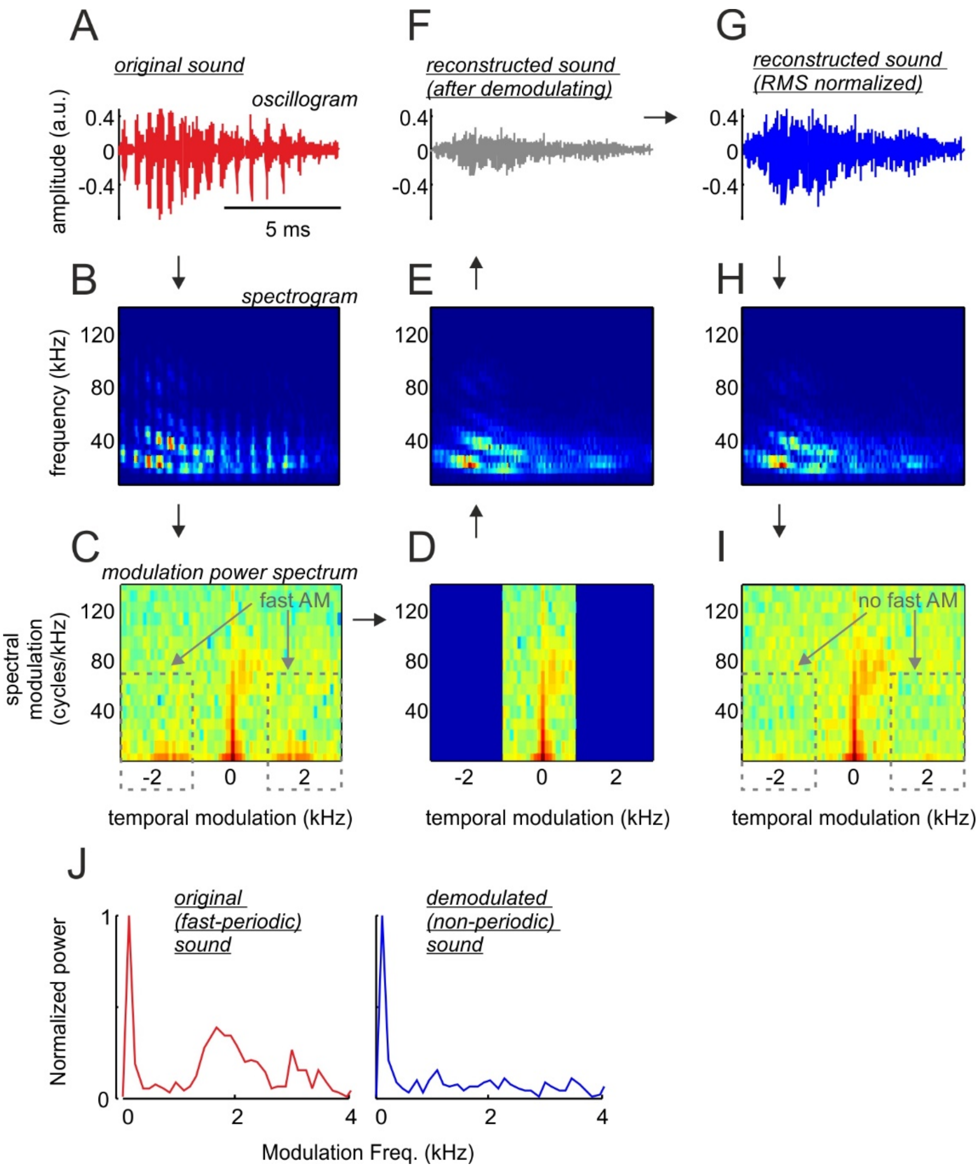


Figure S2. Pipeline used for demodulating fast periodic vocalizations (FPVs). The spectrogram (**A**), waveform (**B**), and modulation power spectrum (MPS, **C**) of an example FPV are shown. **D**: In the MPS, the power at modulation frequencies above 1 kHz and below -1 kHz was set to zero (blue areas in D). **E**: The short time Fourier transform (STFT) of the demodulated vocalization was constructed from the filtered MPS using an inverse FFT2 procedure. **F**: The waveform of the demodulated vocalization was obtained from the STFT represented in E using the weighted overlap add method (WOLA). **G**: The demodulated sound obtained was root-mean-square (RMS) normalized to match the energy of the natural FPV represented in A. **H**: Because the WOLA method can be affected by spectro-temporal trade-offs, the STFT of the demodulated sound shown in G was computed and compared to the "desired" STFT shown in E (see methods). The error was calculated as the squared difference between the desired STFT (shown in E) and the observed STFT (shown in H) divided by the values in the desired STFT. The resulting error value in this case was 1%. **I**: the MPS of the demodulated sound. **J**: The temporal modulation spectra of the natural FPV (left) and its demodulated treatment (right).

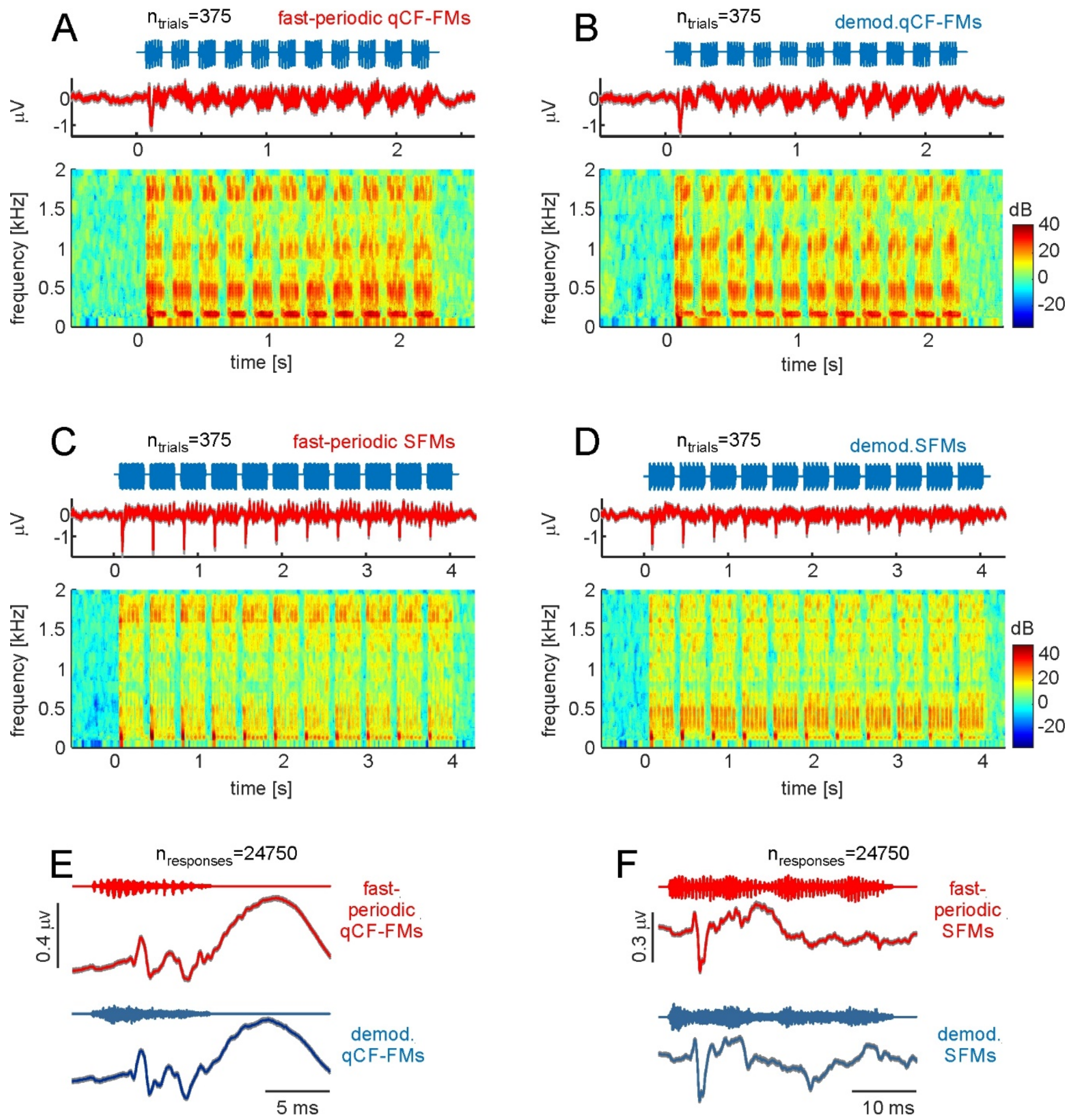


Figure S3. Neural responses to fast periodic and demodulated syllables. (**A** and **B**) Neural responses to a sequence of fast periodic qCF-FM syllables (A) and demodulated qCF-FM (B). Responses are represented as voltage vs. time and in the form of neural spectrograms. (**C** and **D**) Same as A-B but in response to sequences of fast periodic and demodulated SFMs. (**E**) Voltage fluctuations obtained after averaging neural responses to each fast-periodic qCF-FM (red) and each demodulated qCF-FM syllable (blue) across trials and animals. (**F**) Same as E but in response to SFM syllables.
